# Context-specific control of the neural dynamics of temporal attention by the human cerebellum

**DOI:** 10.1101/2019.12.12.874677

**Authors:** Assaf Breska, Richard B. Ivry

**Author notes:** Corresponding author: Assaf Breska, 2121 Berkeley Way, Berkeley, CA 94720-1650.

## Abstract

Physiological methods have identified a number of signatures of temporal prediction, a core component of attention. While the underlying neural dynamics have been linked to activity within cortico-striatal networks, recent work has shown that the behavioral benefits of temporal prediction causally rely on the cerebellum. Here we examine the involvement of the human cerebellum in the generation and/or temporal adjustment of anticipatory neural dynamics, measuring scalp electroencephalography in individuals with cerebellar degeneration. When the temporal prediction relied on an interval representation, duration-dependent adjustments were impaired in the cerebellar group compared to matched controls. This impairment was evident in ramping activity, beta-band power, and phase locking of delta-band activity. Remarkably, these same neural adjustments were preserved when the prediction relied on a rhythmic stream. Thus, the cerebellum has a context-specific causal role in the adjustment of anticipatory neural dynamics of temporal prediction, providing the requisite modulation to optimize behavior.

## Introduction

Temporal anticipation is essential for survival in our dynamic world. Whether playing sports, listening to music, or driving, our brain is able to use temporal regularities to predict the timing of upcoming events [1,2]. These predictions guide proactive allocation of attentional resources and preparation of adaptive responses, expressed in various contexts by the behavioral benefits observed when predictions are confirmed and costs when violated [3–10].

Converging evidence from neurophysiological studies in humans and non-human primates (NHP) have associated temporal prediction and attention with a set of modulations in neural activity in cortico-striatal networks [1]. First, ramping neuronal activity, expressed in human scalp recordings as the contingent negative variation (CNV) potential, is adjusted such that it peaks near the expected time of an upcoming event [6–8,11–14]. Second, on the output end, the power of movement-related beta-band activity decreases just prior to the expected time of the imperative [7,15–20]. Third, an increase in phase consistency of low frequency activity (e.g., delta range, 0.5-3 Hz) is observed across repeated instances of the same interval in temporally predictable contexts, putatively reflecting alignment of high excitability states at an expected time [3,4,6,21,22]. These patterns are evident in NHP recordings in frontal, parietal, and basal ganglia circuits, regions that feature prominently in human fMRI studies of temporal anticipation [9,23–27].

However, we have recently provided evidence that stands in contrast to a strict cortico-striatal view of temporal anticipation, showing that individuals with cerebellar degeneration (CD) failed to exhibit behavioral benefits from temporal cues on a simple detection task [28]. Importantly, the impairment was limited to conditions in which temporal anticipation was based on associations between cues and isolated intervals [6,8–10], but not when based on a periodic signal [3–7]. These findings established a context-specific causal role of the human cerebellum in temporal anticipation.

A fundamental question concerns the role of the human cerebellum in the neural dynamics of temporal anticipation. Electrophysiological recordings in NHP have shown beta-band activity and ramping activity in the cerebellum during timed movements [29–32], and neuroimaging studies have reported increased cerebro-cerebellar connectivity during temporal anticipation [33,34]. However, it is not clear whether the cerebellar activity provides a modulatory input to cortico-striatal circuits involved in temporal prediction, reflects cortico-striatal activity projected to the cerebellum, or occurs independently of cortico-striatal activity.

To address this question, we measured electroencephalography (EEG) in individuals with cerebellar degeneration (CD) while they performed temporal prediction tasks, using either interval-based or rhythm-based cues. Based on our previous study [28], we expected a selective behavioral impairment on the former in the CD group. Our first goal was to examine the causal role of the cerebellum in the CNV, beta-band amplitude and delta-band phase locking when prediction is based on an interval representation. We considered three models (Figure 1A). First, the cerebellum may be critical for the generation of extracerebellar (e.g., cortico-striatal) ramping, delta-band, and/or beta-band activity. By this “cerebellar generation” model, these patterns would be abolished or severely attenuated in the CD group. Second, the cerebellum may be critical for the temporal adjustment, rather than generation, of these neural patterns. By this “cerebellar adjustment” model, the CD group would still demonstrate these neural patterns prior to anticipated events, but no duration-dependent adjustments would be observed in the CNV and beta suppression, and delta phase-locking would not increase to a similar extent as in controls. Third, the cortico-striatal mechanisms reflected in these patterns may operate independently of the cerebellum. By this “cerebellar independent” model, EEG patterns in the CD group would be similar to those observed in control participants.

**Figure 1.**
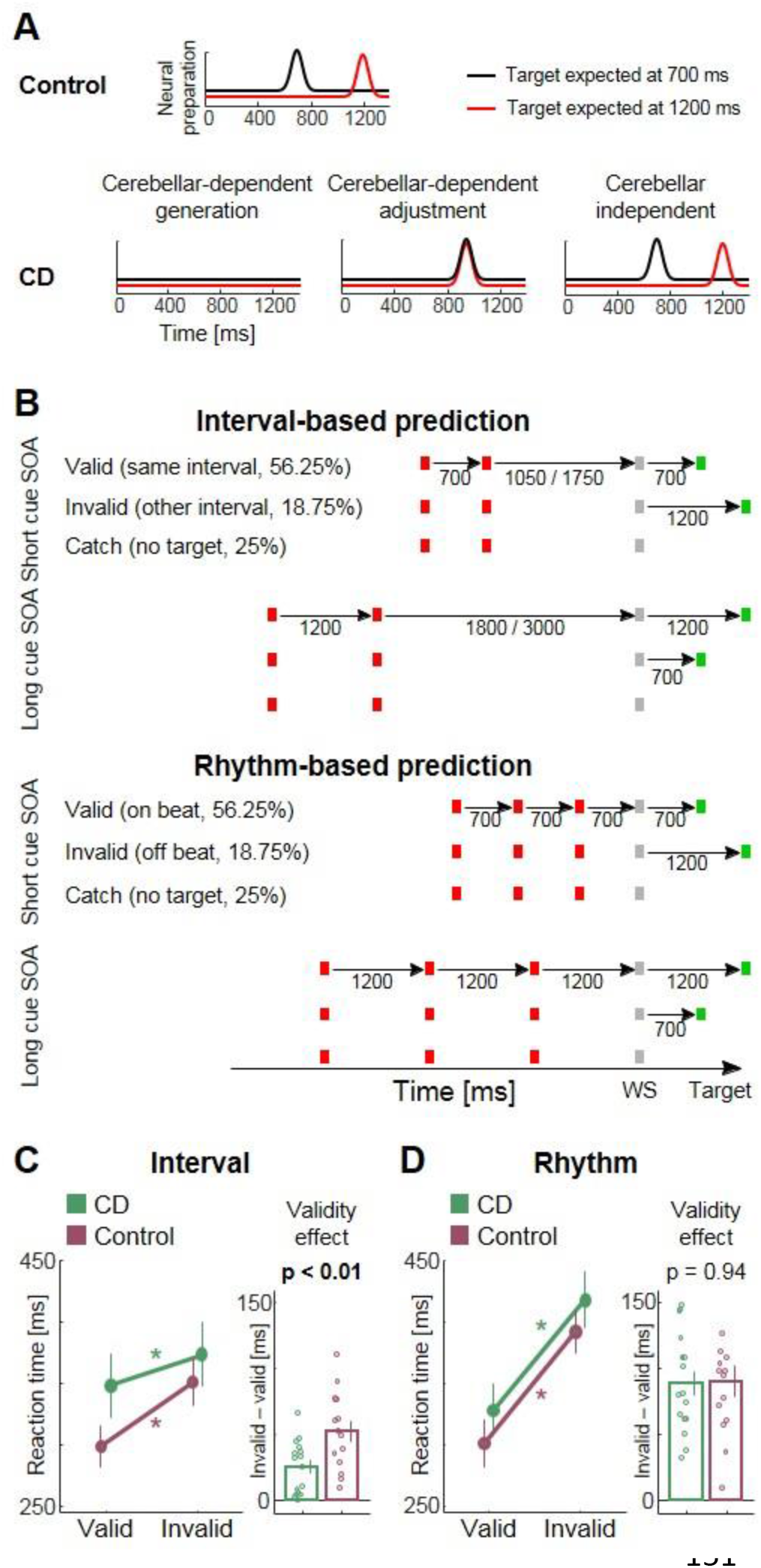
Paradigm and behavioral results. **A.** Possible models of the cerebellar role in neural preparation (arbitrary units). Top: Controls are expected to show adjustment of neural preparation according to expected interval. Bottom: Predictions for CD group for three models. Left: cerebellar-dependent generation of neural preparation predicts no neural preparation in CD group. Middle: Cerebellar-dependent adjustment predicts neural preparation that is not adjusted by expected interval. Right: Cerebellar-independent predicts similar pattern as controls. **B.** Experimental paradigm. Participants viewed a stream of flickering colored squares and provided a speeded response upon detecting a target (green square). Top: Interval task. Two red squares were separated by either a short or long SOA. After a variable delay period, a warning signal (white square) appeared, preceding the target. The interval between the WS and target could be the same SOA (valid) or the other SOA (invalid). Bottom: Rhythm task. Three red squares and WS appeared with identical SOA (short/long). Valid and invalid trials were as in the Interval task. In both tasks, short SOA = 700 ms, long SOA = 1200 ms. On 25% of the trials, no target was presented (catch trials), and participants were to withhold response. **C.** Behavioral results. Line graphs (left) depict mean reaction times (RTs) for the cerebellar degeneration (CD) and control groups for the Interval task. Bar graphs (right) depict magnitude of validity effect (Invalid – Valid). The validity effect is smaller in CD group. **D.** Same as C for the Rhythm task. The validity effect has similar magnitude in both groups. * *p* < 0.05. In both C and D, error bars represent one standard error of the mean (SEM).

A second goal was to address the current debate in the literature regarding the context specificity of these neural signatures of temporal prediction [35–39]. Increased delta-band phase locking has been interpreted as reflecting rhythm-specific prediction mechanisms, such as the entrainment of endogenous oscillations [3,4,21]. However, similar neural adjustments are observed for aperiodic streams that enable interval-based prediction [6]. This has motivated the hypothesis that rhythmic predictions may be mediated by the repeated operation of an interval-based mechanism given that a periodic stream consists of a series of concatenated intervals. Alternatively, similar neural adjustments in rhythm- and interval-based contexts may result from common downstream processes that are driven by context-specific mechanisms. Our previous study [29] provided behavioral evidence consistent with the latter hypothesis, with different subcortical pathologies producing dissociable context-specific impairments. It remains to be seen if similar dissociations are evident in terms of these neural signatures.

Comparing the EEG patterns of interval- and rhythm-based prediction within the CD group provides a novel way to address this issue. Behaviorally, we expected to replicate our previous dissociation, observing a selective deficit in the interval-based condition, with performance similar to matched controls on the rhythm-based condition [28]. For each neural pattern that is impaired in interval-based prediction, finding a comparable impairment in rhythm-based prediction would suggest that it is driven by cerebellar-dependent interval-based mechanisms, even in rhythmic contexts. In contrast, finding a selective impairment in the interval context would suggest that the neural pattern is modulated by an upstream, context-specific mechanism such as a cerebellar-dependent representation of interval information.

## Results

Individuals with cerebellar degeneration (CD, n=16) and controls (n=14) completed two variants of a temporal prediction task in the visual modality [6,28] (Figure 1B). The participants performed a detection task, making a button press as quickly as possible in response to the presentation of a target that appeared after a warning signal (WS). The WS was preceded by a cue that indicated the likely interval between the WS onset and target onset (stimulus onset asynchrony, SOA). In the Interval task, the cue consisted of two stimuli whose SOA was either 700 or 1200 ms. The WS appeared after an interval of either 1.5 or 2.5 times the cue SOA, making the stimulus stream non-isochronous. In the Rhythm task, the temporal cue was defined by a periodic stream of four stimuli, with the last stimulus the WS. The stimuli were separated by a fixed SOA of 700 or 1200 ms. In both tasks, the SOA between the WS and target matched the cue SOA on 75% of the trials (valid trials); on 25%, the WS-target interval matched the other SOA (invalid trials). We also included catch trials in which no target appeared, to discourage premature responses [10].

### Reduced behavioral benefits of interval-based, but not rhythm-based temporal prediction in cerebellar degeneration

Temporal anticipation typically leads to facilitated behavioral performance at expected times, relative to unexpected times or a temporally random sequence [5,8,9]. In our design, this would be manifest as slower reaction times (RTs) on invalid trials compared to valid trials (validity effect). In the Interval task (Figure 1C; see Figure S1 for single-subject data), there was a significant validity effect across groups (mixed analysis of variance [ANOVA], main effect of Validity factor: *F*_(1,28)_ = 72.7, *p* = 2E-9, 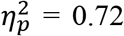). Planned contrasted revealed a significant validity effect in both the CD and control groups (repeated-measures ANOVA, Control: *F*_(1,13)_ = 43.7, *p* = 1E-5, 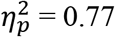; CD: *F*_(1,15)_ = 25.9, *p* = 1E-4, 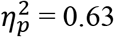). Crucially, the magnitude of the validity effect was significantly smaller in the CD group (25 ms and 53 ms for the CD and control groups, respectively; Validity X Group interaction, *F*_(1,28)_ = 8.75, *p* = 0.006, 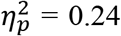). Across groups, the validity effect was also larger for early versus late targets (Validity X Target SOA interaction: *F*_(1,28)_ = 4.96, *p* = 0.034, 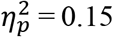), consistent with previous studies using similar tasks [8,10].

A different pattern was observed on the Rhythm task (Figure 1D). As with the Interval task, the validity effect was significant across groups (*F*_(1,28)_ = 138.9, p = 2E-12, 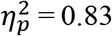), as well as within each group (Control: *F*_(1,13)_ = 54.9, p = 3E-6, 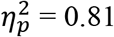; CD: *F*_(1,15)_ = 89, p = 1E-7, 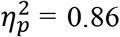). However, there was no significant difference between groups in the magnitude of the validity effect (CD: 89 ms, Control: 91 ms; Group x Validity interaction, *F*_(1,28)_ = 0.006, p = 0.94, 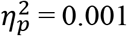; Bayes factor: *B_01_* = 2.9, weak evidence in favor of the null hypothesis). Across groups, there was also a significant main effect of Target SOA, with faster RTs for early versus late targets (*F*_(1,28)_ = 23.4, p = 5E-5, 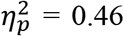), and a smaller validity effect for early versus late targets (Validity X Target SOA interaction: *F*_(1,28)_ = 7.96, p = 0.01, 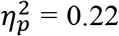). Within the CD group, a comparison of the two tasks revealed a larger validity effect in the Rhythm task (Validity X Task interaction: *F*_(1,15)_ = 54.9, p = 2E-6, 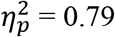).

In sum, the behavioral data show that the participants were faster to detect a target that appeared at a cued point in time, compared to when the target appeared at an unexpected time. However, the CD group exhibited a reduced validity effect for interval-based predictions, but a normal validity effect for rhythm-based predictions.

### Cerebellar degeneration abolishes temporal adjustment of CNV buildup in interval-based prediction

Temporal anticipation leads to adjustment in the buildup of anticipatory ramping neural activity as a function of the duration of the expected interval, such that it peaks just prior to the expected time [11–14]. In human EEG, this is expressed in the CNV, slow ramping in negative potential typically elicited following a WS [6–8]. To test the role of the cerebellum in this temporal modulation, we analyzed the CNV in a pre-defined window prior to the early target time, where the difference between expecting the target at the early and late times should be maximal. The CNV should be more negative amplitude following the short cue relative to the long cue. We also analyzed the CNV across the whole interval between the WS and early target time, without restriction to a specific time range, using a cluster-based permutation analysis [40].

Across groups and tasks, a clear CNV was elicited after the WS with a typical fronto-central scalp distribution (Figure 2A). In the Interval task (Figure 2B), the controls showed a CNV buildup, expressed as a significant difference from baseline, following both cue durations (short: *t*_(13)_ = −4.7, p = 0.0004, Cohen’s *d* = 1.26; long: *t*_(13)_ = −2.67, p = 0.019, Cohen’s *d* = 0.71). Indicative of their sensitivity to the temporal cue, the CNV amplitude was more negative in the predefined window for the short relative to long cue durations (pre-defined window: *t*_(13)_ = −3.12, p = 0.008, Cohen’s *d* = 0.83; cluster-based permutation *p* < 0.05), similar to previous observations in younger individuals [6–8]. The CD patients also showed a CNV buildup for both cues (short: *t*_(15)_ = −4.54, p = 0.0004, Cohen’s *d* = 1.14; long: *t*_(15)_ = −6.65, p = 8E-6, Cohen’s *d* = 1.66). However, the CNV amplitude did not differ between the two cue durations (*t*_(15)_ = 0.1, p = 0.92, Cohen’s *d* = 0.03; Bayes factor: *B_01_* = 3.9, moderate evidence in favor of the null hypothesis; strongest cluster *p* > 0.5). A direct comparison revealed that the modulation of the CNV amplitude by cue duration was significantly larger in controls (Group x Cue SOA interaction: *F*_(1,28)_ = 4.33, p = 0.046, 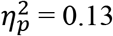). Thus, when temporal prediction relied on timing cued by an isolated interval, the CD group failed to exhibit temporal adjustment in the buildup of the CNV.

**Figure 2.**
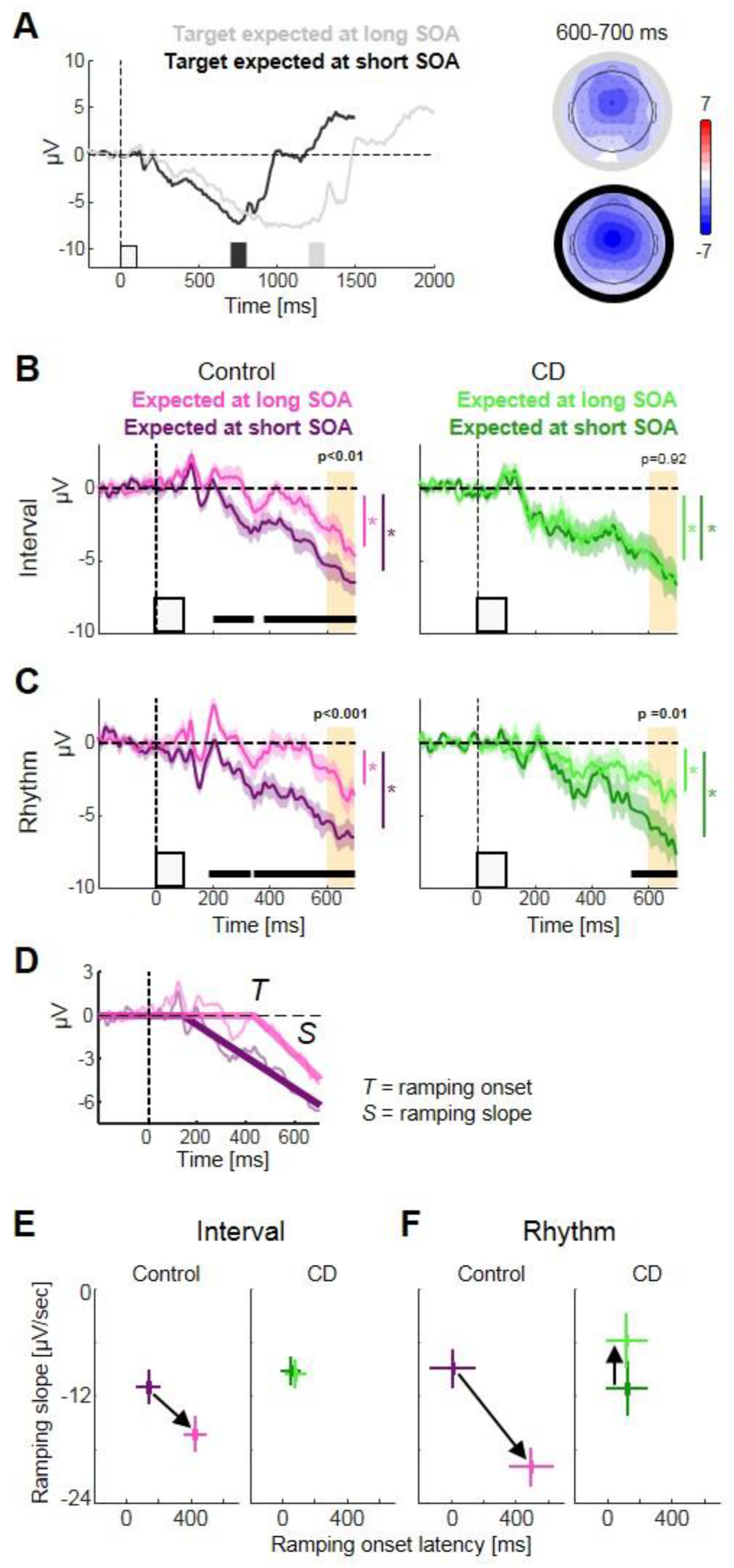
CNV adjustment depends on the cerebellum in interval-based but not rhythm-based prediction. A. CNV elicited following the onset of the warning signal (0 ms) for the short and long cue SOA conditions, averaged across groups and tasks. Black and gray squares indicate expected target onset times. Scalp distribution depicted for a time window preceding the short SOA target (black and gray outlines correspond to short and long conditions). **B.** CNV in the Interval task for the two groups. Yellow background indicates the pre-defined window for analysis of the CNV amplitude as a function of the expected interval. Horizontal bars indicate significant clusters of consecutive time points showing difference between conditions (*p* < 0.05). Differential CNV buildup is observed in Control group (left) but not in the CD group (right). Error margins indicate one SEM of the difference between expected SOAs. (\**p* < 0.05 against baseline). **C.** Same as B for the Rhythm task. Differential CNV buildup is observed in both groups. **D.** Ramping activity model. The CNV buildup was fitted with a two-parameter segmented regression model, allowing variation of ramping onset time *(T)* and ramping slope (*S*). The depicted model fit is for the control data in the Interval task. **E.** Parameter estimates of the ramping activity model in the Interval task. Black arrow links estimate from short SOA condition to long SOA condition. Controls show delayed ramping onset and steeper slope in the long SOA condition. There was no difference between the two SOA conditions for the CD group. Horizontal and vertical error bars indicate one SEM of the difference between expected SOAs for ramping onset latency and slope, respectively. **F.** Same as E for the Rhythm task. The control group showed a similar pattern as in the Interval task, whereas the CD group showed a difference between two SOA conditions, manifest only as slope modulation, with a lower slope in the long SOA condition.

In the Rhythm task, both groups showed reliable CNV buildup for both cue durations (Short: controls: *t*_(13)_ = −5.41, p = 0.0001, Cohen’s *d* = 1.45; CD: *t*_(15)_ = −5.1, p = 0.0001, Cohen’s *d* = 1.28; Long: controls: *t*_(13)_ = −3.95, p = 0.0017, Cohen’s *d* = 1.06; CD: *t*_(15)_ = −3.63, p = 0.0025, Cohen’s *d* = 0.91; Figure 2C). Similar to the Interval task, the CNV for the controls was more negative in the predefined window in the short cue durations (predefined window: *t*_(13)_ = −4.44, p = 0.0007, Cohen’s *d* = 1.19; cluster-based permutation analysis, *p* < 0.05). In contrast to the Interval task, the CD group also showed a more negative CNV in the short cue duration (*t*_(15)_ = – 2.96, p = 0.01, Cohen’s *d* = 0.74). Consistent with this, the permutation analysis also revealed a significant cluster of time points in which the CNV was more negative in the short SOA condition (*p* < 0.05), although with a shorter temporal extent than in the control group (see next section). Direct comparison between groups revealed no difference in the magnitude of CNV modulation by the cue duration between the two groups (Group X Cue SOA interaction: *F*_(1,28)_ = 0.003, p = 0.96, 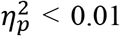; Bayes factor: *B_01_* = 3.02, moderate evidence in favor of the null hypothesis). Finally, the direct comparison between the two tasks within the CD group showed a larger effect of cue duration in the Rhythm task (Cue SOA X Task interaction: *F*_(1,15)_ = 6.78, p = 0.02, 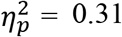).

Thus, when temporal prediction relied on timing cued by an isolated interval, the CD group failed to exhibit temporal adjustment in the buildup of the CNV. In contrast, when temporal predictions relied on a rhythm, the CD group showed an adjustment of the CNV buildup, leading to similar difference in CNV amplitude prior to the early target SOA as in controls.

### CNV modulation in the Rhythm task for CD group is manifest as change in slope rather than latency adjustment

Given that the CNV is reflective of ramping activity, its shape can be informative concerning the dynamics of preparatory resource allocation. Adjustment of ramping activity in response to different temporal goals can involve modulation of either its onset latency or its slope, as evident in NHP neural recordings [12,31,32,41]. We fit the CNV waveform generated between the WS onset and the earliest possible target time (700 ms) with a two-parameter linear ramping model (Figure 2D), one corresponding to the slope and the other with onset latency. Four fits were performed for each group (two tasks and two cue SOAs), with the data averaged across participants within a group.

In the Interval task, the modeling results showed a difference between the two cue durations for both parameters for the control group (Figure 2E, permutation test, both *p’s* < 0.05): The latency was earlier and slope shallower following the short cue compared to the long cue. In contrast, neither parameter differed for the CD group (both *p’s* > 0.5). A direct comparison revealed a significant Group x Cue SOA interaction for both latency (*p* < 0.05) and slope (*p* < 0.05). Thus, these results indicate that temporal modulation of ramping activity, either via latency or slope, was impaired and even absent in the CD group.

In the Rhythm task (Figure 2F), the controls showed a parameter difference in latency (*p* < 0.05), and slope (*p* < 0.05) between the two cue durations, similar to what was observed in the Interval task. For the CD group, there was no difference in the parameter estimates of latency (*p* > 0.5). However, there was a significant difference in slope (*p* < 0.05), with the slope steeper following short duration cues. Direct comparison revealed a significant Group x Cue SOA interaction for both parameters (*p’s* < 0.05). When directly comparing the two tasks, a larger adjustment of the slope parameter was observed in the Rhythm task compared to the Interval task in the CD group (*p* < 0.05), with no difference in the adjustment of the latency parameter by the expected interval (*p* > 0.05). Note that the slope difference for the CD group on the Rhythm task is in opposite direction than that found for the controls. When a rhythmic cue indicates a longer interval, the CNV is delayed in controls and then builds up at a faster rate. In contrast, for the CD group, the latency remains unchanged but the slope is shallower, leading to a slower buildup.

### Cerebellar degeneration reduces temporal alignment of low-frequency activity in interval-based but not rhythm-based prediction

Temporally predictive streams are associated with phase alignment of low-frequency activity during the anticipatory period [3,4,6]. This is expressed as an increase in phase consistency across trials for a given expected target interval, and typically quantified using the inter-trial phase concentration (ITPC) index [42]. To test the dependence of phase alignment on the cerebellum, we band-pass filtered the EEG data to a delta-band frequency range that corresponds to the expected intervals (0.6-2 Hz, see Methods), and calculated the ITPC following the WS (Figure S2). We focused on EEG data recorded in fronto-central electrodes, using the same electrodes as in the CNV analysis.

We first examined whether the ITPC differed between the two groups in the absence of temporal prediction, in a baseline period of 100 ms prior to the onset of the warning signal in the Interval task (Figure 3B). Note that it is not possible to extract a comparable baseline for the Rhythm task given that the onset of the WS is also predictable. During the baseline epoch, there was no difference in ITPC between the control and CD groups (*p* > 0.3, between-subject permutation test, Figure 3C). In the Interval task, temporal anticipation led to increased ITPC just prior to the time of the target relative to baseline in both groups (600-700 ms after the WS, CD: *p* < 0.01; Controls: *p* < 0.001, within-subject permutation test, Figure 3B). However, the ITPC was higher in the control group (*p* < 0.05, Figure 3C; see Figure S2 for single-subject data). Thus, the increase in ITPC of low-frequency activity in interval-based prediction depends on the cerebellum.

**Figure 3.**
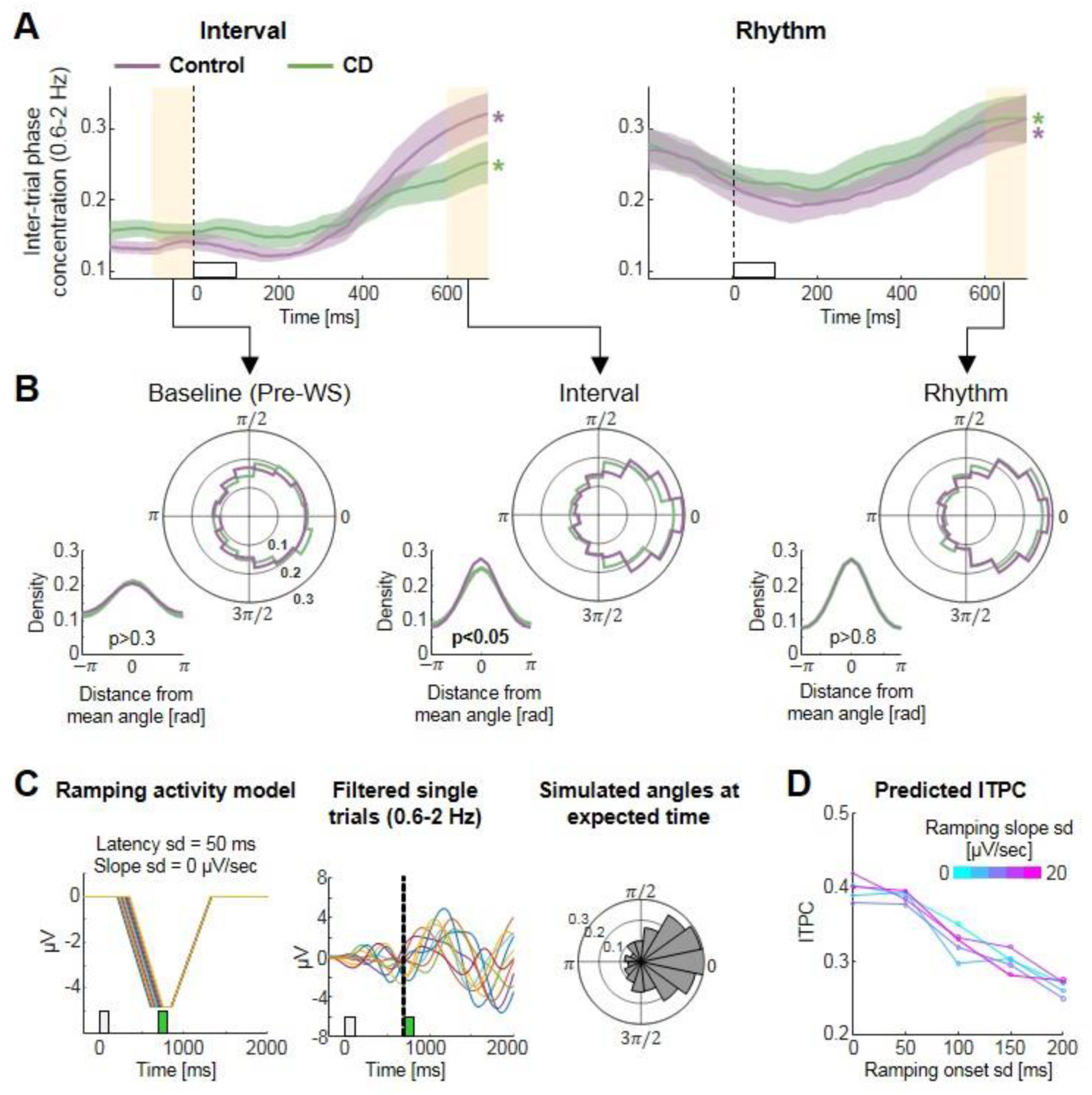
Temporal alignment of neural activity depends on the cerebellum in interval-based, but not rhythm-based prediction. **A.** Time-series of inter-trial phase concentration (ITPC) in the delta frequency range (0.6-2 Hz), locked to the WS onset (white square) for the Interval task (left) and Rhythm task (right). Target anticipation is associated with gradual ITPC increase from baseline in both groups on both tasks. Error margins indicate one SEM of the difference between groups. Yellow background indicates the baseline period (left) and pre-target window for analysis (right). * *p* < 0.05 for difference against baseline. **B.** Phase distributions of delta-filtered neural activity in the baseline phase (left), and pre-target window for the Interval (middle) and Rhythm (right) tasks. The control group exhibits stronger ITPC than the CD group in the Interval task only. **C.** Modelling ITPC in the Interval task. Left: Simulated ramping activity elicited following the WS (white square), with return to baseline after target onset (green square), for a set of trials with no variability in slope and low variability in onset latency. Middle: Same simulated trials, band-pass filtered to the delta range (0.6-2 Hz). Right: Circular distribution of phase angles just prior to target time in the simulated trial set. **D.** Predicted ITPC levels in simulated trial sets as function of the inter-trial variability in ramping onset latency and in ramping slope. ITPC depends on the magnitude of variability in ramping onset. sd = standard deviation.

An ongoing debate concerns whether the finding of similar ITPC in rhythm- and intmediated by a cerebellar-dependenterval-based prediction in neurotypical participants [6] reflects different context-dependent temporal prediction mechanisms, or a single mechanism that operates in both contexts. For example, predictions in rhythmic streams could be mediated by rhythm-specific mechanisms, such as oscillatory entrainment, or by repeated interval-based processes [6,35–37]. If the latter was true, the CD group should also show reduced ITPC relative to controls in the Rhythm task, reflecting their impairment in the Interval task. However, the ITPC values were similar for the control and CD groups (*p* > 0.8, Figure 3C; see Figure S2 for single-subject data; ITPC increase relative to baseline CD: *p* < 0.001; Controls: *p* < 0.001). Furthermore, direct comparison of the Rhythm and Interval tasks in the CD group revealed significantly stronger ITPC in the Rhythm task (*p* < 0.01, within-subject permutation test). Thus, the ITPC increase in rhythm-based prediction does not rely on cerebellar-dependent interval mechanisms.

The selective impairment in the Interval task indicates that the ITPC increase observed in the control group (as well as in other studies [6]) is mediated by a cerebellar-dependent interval-based mechanism. To explore this hypothesis, we used a modeling approach to ask how an increase in ITPC might arise from a non-oscillatory mechanism. Given that ITPC is essentially a measure of variability, we examined the impact of temporal variability in ramping activity on ITPC. We simulated a set of trials with non-oscillatory ramping activity using the 2-parameter ramping model described above, adding the constraint that the CNV returns to baseline after target onset. These data were simulated for a condition in which the cue indicated an early target (700 ms). We then applied the same ITPC analysis used with the EEG data (Figure 3C). We repeated this simulation manipulating the inter-trial variability of ramping onset latency or ramping slope, asking how these manipulations impact ITPC. This analysis revealed that ITPC decreases with larger inter-trial variability in ramping onset latency (*t*_(74)_ = −6.2, p = 3E-8), but is not affected by larger inter-trial variability in ramping slope (*t*_(74)_ = −1.04, p =, 0.3, Figure 3C). Thus, countering the idea that ITPC modulations selectively reflect oscillatory alignment, our analyses here show that a reduction in ITPC can also reflect reduced temporal consistency of ramping activity. We conjecture that this might underlie the pattern observed in the CD group.

### Timed suppression of beta activity is impaired in cerebellar degeneration in interval-based but not rhythm-based prediction

We next analyzed the amplitude of beta-band activity, a signal associated with the transition from motor preparation to movement initiation. Temporal anticipation is associated with amplitude decrease of non-phase-locked beta-band activity prior to a response [7,15–17]. We tested the dependence of this on the cerebellum by conducting a time-frequency analysis, asking whether expecting a target (and, as such, the subsequent response) at the early interval would lead to decreased beta amplitude compared to when expecting the target at the later interval. To avoid contamination from target-evoked activity, we only analyzed data from trials in which the target either appeared at the long SOA or from catch trials [6].

Consistent with the literature, beta suppression was prominent in central-parietal electrodes (see Figure S3). In the Interval task, beta amplitude decreased in both groups following presentation of the WS (Figure 4A). We first analyzed beta-band activity in a time window just prior to the time of the early target (500-700 ms post-WS, 14-26 Hz). In the control group, beta band amplitude was relatively reduced following short cues compared to long cues (*t*_(13)_ = −2.7, *p* = 0.018, Cohen’s *d* = 0.72, Figure 4B). In contrast, in the CD group no relative reduction in beta amplitude was observed (*t*_(15)_ = −0.15, *p* = 0.88, Cohen’s *d* = 0.04, Bayes factor: *B_01_* = 3.88, moderate evidence in favor of the null hypothesis). Direct comparison of the two groups revealed significantly stronger modulation of beta amplitude in the Control group (Group x Cue SOA interaction: *F*_(1,28)_ = 4.46, *p* = 0.04, 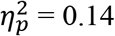).

**Figure 4.**
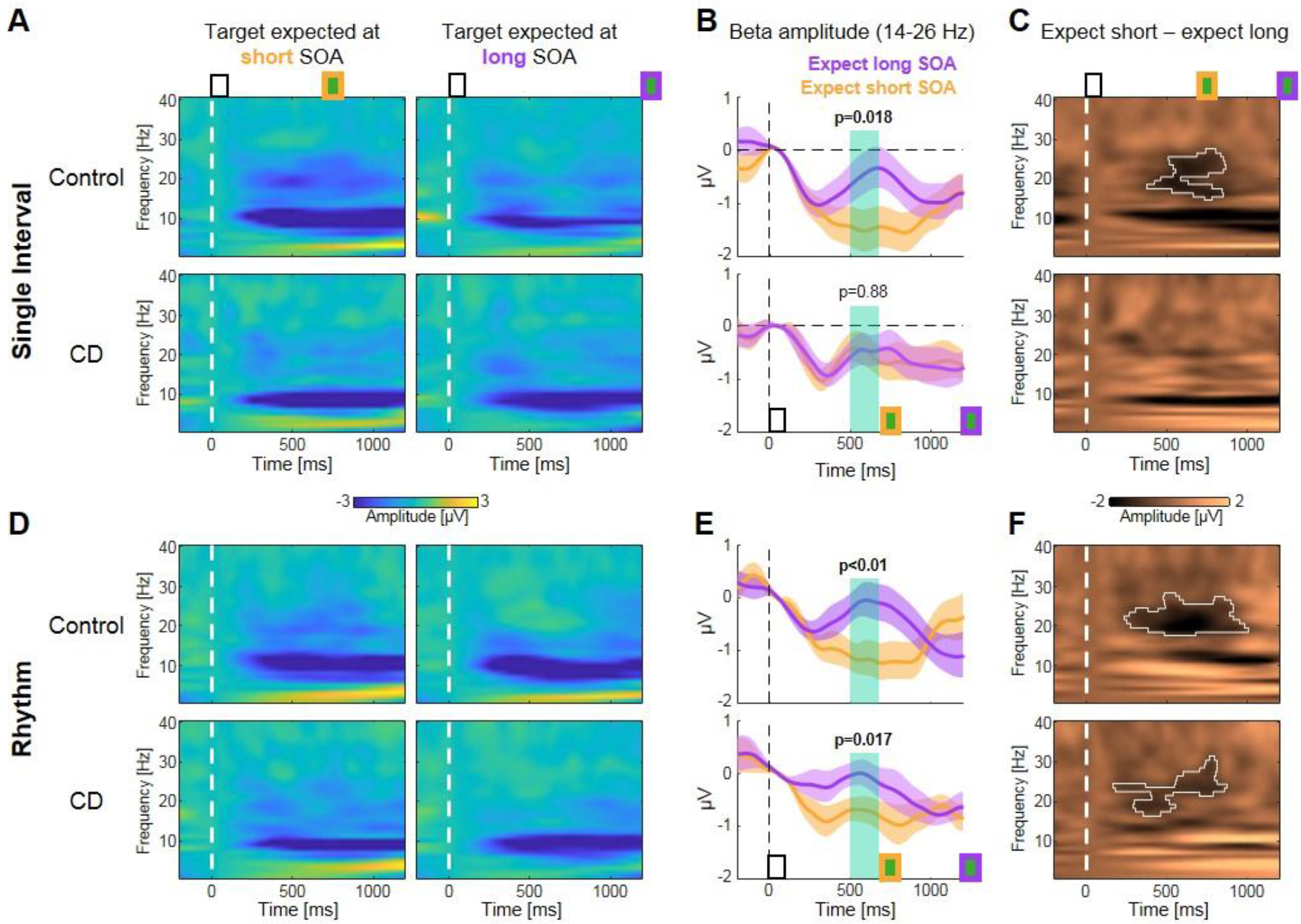
Timed suppression of beta-band activity depends on the cerebellum in interval-based but not rhythm-based prediction. **A.** Time-frequency amplitude representations in the Interval task for the control (top) and CD groups (bottom), referenced to a pre-WS baseline period. Left: short cue SOA, target omitted. Right: long cue SOA. Vertical dashed line indicates WS onset. Both groups show amplitude decrease in a broad frequency range. **B.** Beta amplitude (14-26 Hz) dynamics in the Interval task for the control (top) and CD groups (bottom). Around the short SOA target time, beta amplitude is decreased following short cue SOA in the control group, but not in the CD group. Error margins indicate one SEM of the difference between cue SOAs. Green background indicates pre-defined pre-target window for analysis. **C.** Time-frequency representations of the amplitude difference between expecting the target at short and long SOAs. A significant cluster in the beta range is only found for the controls (outlined in white, p<0.05). Vertical dashed line indicates WS onset. **D-F.** Same as A-C for the Rhythm task. Both groups show amplitude decrease in the beta range following short relative to long SOA cues.

In addition to analyzing amplitude in a pre-defined time-frequency window, we also compared the conditions using a two-dimensional cluster-based permutation approach (Figure 4C). Whereas a significant cluster of reduced beta amplitude when expecting the target at a short versus long interval was observed in the control group in the beta range (*p* < 0.05), no significant clusters were found in the CD group (strongest cluster *p* > 0.5). Direct comparison between groups confirmed that the controls showed a stronger reduction in beta amplitude (*p* < 0.05). Thus, both analyses converge, indicating that cerebellar degeneration impairs the anticipatory temporal adjustment of beta-band suppression when temporal anticipation is based on an interval representation.

A different picture emerged in the analyses of the data from Rhythm task. Both groups again showed beta amplitude decrease following presentation of the WS (Figure 4D). Analysis in the pre-defined time-frequency window revealed reduction in beta amplitude following the short cue compared to the long cue in both groups (controls: *t*_(13)_ = −3.52, *p* = 0.004, Cohen’s *d* = 0.94; CD: *t*_(15)_ = −2.68, *p* = 0.017, Cohen’s *d* = 0.67, Figure 4E). Direct comparison of the two groups did not show a significant Group x Cue SOA interaction (*F*_(1,28)_ = 1.17, *p* = 0.29, 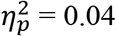). The cluster-based permutation analysis revealed significant clusters of reduced amplitude in the beta range for both groups (*p’s* < 0.05, Figure 4F), and the direct comparison of the effect of cue duration between the two groups showed no significant clusters (strongest cluster *p* > 0.5). Thus, both analyses converge on indicating that cerebellar degeneration did not impair the anticipatory temporal adjustment of beta-band suppression by rhythm-based predictions. Finally, the comparison between tasks in the CD group revealed stronger beta suppression following short compared to long cues in the Rhythm task in the cluster-based permutation test (*p* < 0.05); in the pre-defined window this analysis did not reach significance (*F*_(1,15)_ = 2.14, *p* = 0.16, 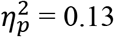).

### Cerebellar degeneration results in reduced target-evoked P3 response in interval-based but not rhythm-based prediction

In the final analysis, we asked whether the cerebellar-dependent impairment in temporal adjustment on the Interval task would translate into target processing as reflected in target evoked activity. We focused on the P3 response, a parietal-maximum positive polarity ERP associated with stimulus evaluation and response selection [43,44]. This response is enhanced when a temporal prediction is confirmed relative to when the stimulus occurs at an unexpected time [8,45]. We analyzed the P3 amplitude in a parietal electrode cluster in response to early targets (appearing at 700 ms), comparing the P3 on valid and invalid trials (see Methods). We used a pre-defined time window of 275-325 ms after the target, as well as a cluster-based permutation approach [40].

Across groups and tasks, the P3 response was enhanced on valid, relative to invalid trials (Figure 5A). In the Interval task (Figure 5B), controls displayed a significant P3 enhancement (*t*_(13)_ = 3.43, *p* = 0.005, Cohen’s *d* = 0.92; cluster-based permutation *p* < 0.005). However, in the CD group analysis in the pre-defined window yielded only a marginally significant effect, and no cluster was detected in the cluster-based approach (*t*_(15)_ = 1.88, *p* = 0.08, Cohen’s *d* = 0.47; strongest cluster *p* > 0.25). Direct comparison between groups revealed a significant Group x Validity interaction (*F*_(1,28)_ = 5.66, *p* = 0.024, 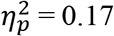), indicating that the enhancement of the P3 was smaller in the CD group.

**Figure 5.**
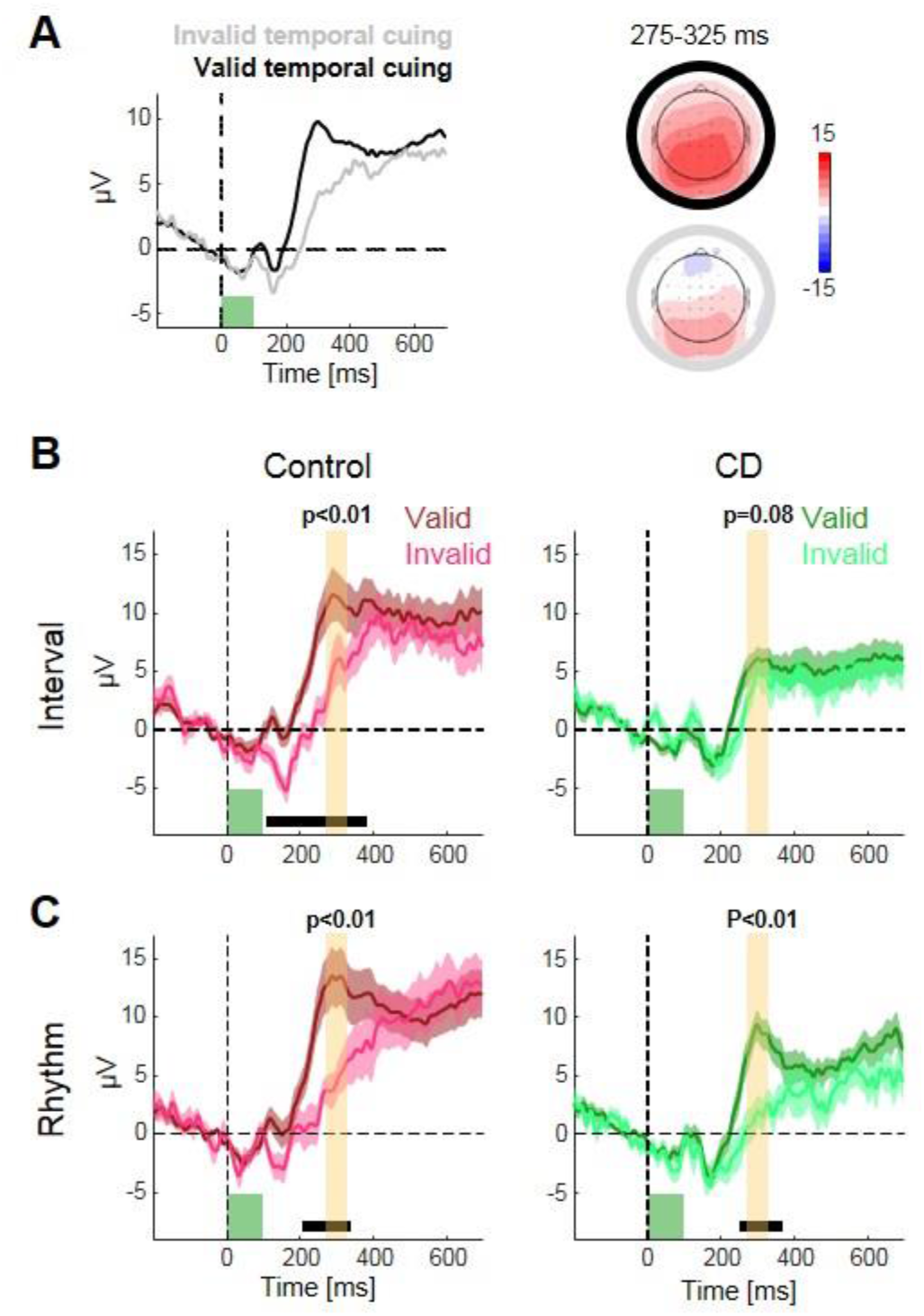
P3 enhancement depends on the cerebellum in interval-based but not rhythm-based prediction. **A.** ERPs time-locked to target onset (green square) on valid (dark) and invalid (light) trials, averaged across groups and tasks. The P3 response is enhanced on valid trials. Scalp distribution depicted for a time window surrounding the P3 peak. **B.** P3 enhancement on valid trials in the Interval task is only observed in the control group (left) but not in the CD group (right). Error margins indicate one SEM of the difference between expected SOAs. Yellow background indicates the pre-defined pre-target window for analysis (* *p* < 0.05). Horizontal bars indicate clusters of consecutive time points with significant difference between conditions (*p* < 0.05). **C.** Same as B for the Rhythm task. P3 enhancement on valid trials is observed in both groups.

In the Rhythm task (Figure 5C), the P3 amplitude was significantly more positive on valid relative to invalid trials for both groups (controls: *t*_(13)_ = 4.2, *p* = 0.001, Cohen’s *d* = 1.12; clusterbased permutation *p* < 0.05; CD: *t*_(15)_ = 3.68, *p* = 0.002, Cohen’s *d* = 0.92; cluster-based permutation *p* < 0.05). The Group x Validity interaction was not significant in a direct comparison between groups (*F*_(1,28)_ = 0.41, *p* = 0.53, 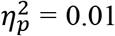). Direct comparison between the two tasks in the CD group revealed that the P3 enhancement was significantly stronger in the Rhythm task compared to the Interval task (Cue Validity X Task interaction: *F*_(1,15)_ = 10.41, p = 0.006, 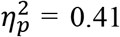). Thus, cerebellar degeneration attenuated the enhancement of P3 from temporal prediction when based on an interval cue, but not when based on a rhythmic cue.

## Discussion

Our results reveal a critical role for the cerebellum in the control of the neural dynamics associated with temporal prediction in an attentional orienting task [1,2]. When temporal predictions were based on an interval-based representation, cerebellar degeneration (CD) impaired several EEG signatures of temporal anticipation, including the CNV, beta-band activity, and phase locking of low-frequency activity. Remarkably, when the expected interval was cued using a periodic stream of stimuli, these same neural signatures were preserved. Echoing this dissociation, the CD group also showed diminished behavioral facilitation as well as an attenuation of the target-evoked P3 response to valid targets in the interval condition, but a normal response in the rhythm condition. These results establish a causal role of the cerebellum in the adjustment of anticipatory neural dynamics by interval-based temporal prediction. Moreover, they indicate that these adjustments of neural dynamics can be driven by distinct mechanisms in different contexts.

### Causal role of the cerebellum in the neural dynamics of temporal prediction when based on isolated interval representation

In line with our previous findings, the CD group showed a reduced behavioral benefit when provided with an interval cue to anticipate the onset of a target stimulus [28], bolstering the claim that interval-based temporal predictions rely on the cerebellum. The primary goal in the current study was to identify how the cerebellum influences prominent neural signatures of anticipation, typically associated with cortico-striatal circuits. As outlined in the Introduction, we considered three models (see Figure 1A). First, these anticipatory signals might operate independently of the cerebellum, perhaps reflecting the ubiquitous nature of preparatory activity. Second, the cerebellum might be the generator of these signals, a hypothesis consistent with cerebellar physiological recordings showing ramping activity and beta-band oscillations during the production of timed movements [29–32]. Third, output from the cerebellum may be essential for the adjustment of these anticipatory signals based on the expected interval.

The current results reveal a critical role of the cerebellum in modulating anticipatory neural dynamics, consistent with the cerebellar adjustment model. We observed CNV buildup, beta activity, and delta ITPC increase in the CD group in response to both types of cues. The fact that these signals were present in both tasks argues against the cerebellar generation model. However, the CD group showed no variation in the CNV buildup or timing of beta-band suppression between the short and long cues in the interval condition, and reduced delta ITPC. Thus, the integrity of the cerebellum is required to modulate anticipatory neural dynamics as a function of the expected interval.

It is important to emphasize that we do not claim that the EEG signals described here directly reflect cerebellar activity. As noted above, evidence from multiple methodologies indicates that these signals reflect activity in cortico-striatal networks [3,7,9,11,19,20,46]. Furthermore, recent work suggests that EEG recordings only capture high frequency cerebellar activity (> 75 Hz, [47]). Thus, abnormalities in the EEG signals in the CD group would presumably reflect the downstream consequences of altered or absent cerebellar input.

In terms of functional interpretation, the CNV and beta activity have been associated with different processing stages. The CNV is observed on both motor and non-motor tasks [48–50]. Consequently, it has been interpreted as reflecting an increase in excitability in preparation for an upcoming event, independent of the response to that event [50,51]. In contrast, suppression of beta-band activity over sensorimotor regions is observed during motor execution and anticipation of movement cues [52,53], suggesting it reflects increased motor readiness [54,55]. Our results imply that when temporal predictions rely on interval representation, the cerebellum is critical to multiple functional aspects of the temporal anticipation process.

### Specificity of cerebellar involvement in temporal anticipation

In striking contrast to their impairments on the Interval task, the CD group exhibited normal behavioral benefits from temporal cueing in the Rhythm task, along with cue-dependent adjustments of the CNV buildup, beta-band suppression, and delta ITPC. The dissociation between the Interval and Rhythm tasks has several important implications. Methodologically, it suggests that the EEG abnormalities in the Interval task in the CD group are unlikely to reflect inferior data quality. Theoretically, the dissociation corroborates the functional separation between interval-based and rhythm-based prediction [28], and specifies an important boundary constraint on cerebellar contributions to prediction and timing. Moreover, it indicates a separation between processes that are involved in the generation of anticipatory neural patterns, and processes that modulate these signals in a context-specific manner.

We recognize that there are inherent limitations in the interpretation of a single dissociation. For example, it is possible that the Rhythm task is less demanding than the Interval task. However, our control group, as well as two previous studies [6,28], showed similar behavioral and neural adjustments in interval and rhythmic conditions. Moreover, we had, a priori, anticipated the observed dissociation: The interval-specific hypothesis arose from our previous behavioral work in which we observed a double dissociation, such that individuals with Parkinson’s disease were selectively impaired in exploiting rhythmic temporal cues ([28], see also [6]). Although we have chosen to focus on the cerebellum in this study, previous work from other labs has shown that PD is associated with impairments in CNV modulation and beta-band attenuation based on rhythm-based predictions [56,57]; future work should examine these measures in response to interval cues. At present, the dissociation within the CD group points to an asymmetric role of the cerebellum in these two modes of prediction, and also implies that the neural signatures of predictions do not rely on the cerebellum across contexts.

Our model-based analysis of the CNV response did reveal one abnormality in the CD group in the Rhythm task: Whereas both onset latency and slope differed between the short and long conditions in the control group, the CD group did not demonstrate an adjustment of onset latency, but only of the slope. Thus, the cerebellum seems to be necessary for latency adjustment of ramping activity across contexts. Consistent with this, neural recordings in non-human primates reveal that, when the delay prior to a cued movement is varied, ramping activity in the cerebellum changes in onset latency but not slope; interestingly, the reverse is observed in the striatum where the slope varies, but not the latency [31,32]. The slope adjustment observed in the CD group in the Rhythm task might reflect greater dependence on striatal computations, essential for rhythmic prediction [29]. This hypothesis can account for the finding that the variation in slope for the short and long cues went in the opposite direction for the CD group compared to the controls. Given their inability to adjust the onset latency, achieving a high state of preparation for shorter intervals requires a faster rise in the CNV.

### Rhythm-specific prediction mechanisms and the role of oscillatory entrainment

Whether temporal prediction relies on oscillatory entrainment is controversial, as is the interpretation of ITPC as a measure of entrainment [35–39]. Increased ITPC of low-frequency activity was originally observed when predictions were formed in periodic streams, and was hypothesized to reflect the phase-alignment of low-frequency oscillations [2–4,21]. However, a similar ITPC increase is observed when aperiodic events enable interval-based predictions [6]. It is not clear if ITPC increases in periodic and aperiodic contexts are driven by distinct mechanisms, or whether a single mechanism explains ITPC modulations in both contexts. For example, ITPC in rhythmic streams could reflect the repeated operation of interval-based mechanisms [6,58].

For the control group, ITPC enhancement was similar in magnitude in the Interval task as in the Rhythm task, despite the fact that the former does not lend itself to oscillatory entrainment. This further confirms that ITPC increase can be explained by interval-based mechanisms [6]. Here, we add to this observation the dissociation in the CD group between the two tasks. Together, these results argue against the hypothesis that ITPC in rhythmic streams reflects the repeated operation of an interval-based mechanism, as this would predict a similar reduction in the Rhythm task. Thus, we propose that an increase in ITPC reflects the operation of different mechanisms when predictions are based on rhythmic streams or an interval-based representation, and that the latter is dependent on the cerebellum. The simulations we present further support this hypothesis, showing that an increase in variability of the onset latencies of non-oscillatory ramping activity, when subjected to band-pass filtering in the delta range, results in a decrease in ITPC. More generally, the modeling provides a demonstration that a change in spectral information need not be reflective of an underlying oscillatory process [59].

Overall, the dissociation provides strong evidence in favor of separate mechanisms for rhythm- and interval-based predictions. However, the current results do not enable concluding whether rhythm-based predictions rely on oscillatory entrainment, or on an alternative mechanism. While the slope adjustment in CD patients in the Rhythm task might seem in line with the idea that oscillators with different frequencies are entrained in the short and long conditions, it is not clear whether this adjustment is specific to an oscillatory mechanism. Strong evidence for involvement of oscillatory mechanisms can only come from observing oscillatory patterns that cannot be explained by interval-based prediction, such as reverberation after stream termination [6,60,61].

### Selective role of the cerebellum in representing isolated interval across timing domains

Identifying the constraints on the role of the cerebellar in temporal processing has been a central goal for research of timing [62,63] and temporal prediction [64,65]. The results from this study indicate that the cerebellum is not essential for attentional anticipation and motor preparation: We observed CNV buildup and beta suppression in both tasks in the CD group. The cerebellum is also not the sole source of temporal adjustments, as these were preserved in the Rhythm task. Other context-independent hypotheses of cerebellar function such as generalized prediction or rapid neural coordination [32,64,66] seem incompatible with the preserved performance observed in the Rhythm task. Instead, cerebellar computation appears to be specific to prediction in an interval-based context, either constituting a unique prediction mechanism for intervals, or providing the representation of the isolated interval in the absence of continuous context or regularity. Given that prediction can also be driven by other sources, as indicated by the preserved rhythm-based prediction, the latter possibility seems more parsimonious.

A selective role for the cerebellum in interval-based timing is also observed in other timing domains [27]. In timed movement, cerebellar degeneration impairs discrete tapping but not circle drawing, where the latter might be achieved by transforming the temporal goal into a velocity-based signal [67]. In explicit timing judgments, cerebellar degeneration impairs interval discrimination but not beat identification [68]. Similar dissociations between isolated interval timing and more continuous forms of timing have been reported in neuroimaging studies [69,70]. Computationally, it has been proposed that the cerebellum is critical for timing isolated intervals that are defined by salient events (‘event timing’), but not when temporal representation is inherent to the ongoing dynamic context (‘emergent timing’) [71]. Our results underscore that this constraint is not limited to tasks involving explicit temporal representation, but also applies to temporal anticipation tasks in which timing is implicit.

Moreover, our results specify the manner in which the cerebellum contributes to temporal anticipation for interval-based predictions. Specifically, the CD group was impaired in two ways with respect to the onset latency of ramping activity. First, they exhibited an impairment in the systematic adjustment of ramping onset based on the expected interval. Second, the reduction in ITPC can be attributed to a reduction in temporal consistency of onset latency across trials (see model-based analysis). Taken together, these impairments indicate that the cerebellum controls interval-based temporal predictions through latency adjustment of anticipatory processes. When intact, this cerebellar-dependent adjustment process would support more cost-effective resource allocation, allowing an appropriately tuned neural state at the expected time.

This hypothesis makes explicit that the neural circuitry underlying temporal prediction entails multiple levels of information processing. Adjustments of the CNV, beta activity and delta ITPC reflect processes related to the control and implementation of attentional and motor preparation [1] that appear to be context-independent. However, to be optimized, these signals require modulatory inputs; in an interval context, the cerebellum provides the context-specific representations of time [72]. In other contexts, such as rhythmic streams, the representation of time, as well as manner in which neural dynamics for temporal anticipation are modulated might rely on other structures such as the striatum [28,73]. Consistent with this hypothesis, temporal adjustment of the CNV and beta activity in these contexts is not impaired by cerebellar dysfunction, but by striatal dysfunction [57,74].

To conclude, our results indicate that the neural dynamics of attentional anticipation in the time domain critically relies on the cerebellum in a context-specific manner, one in which prediction arises from an interval-based representation. The findings not only advance our understanding of the functional domain of the cerebellum in timing, but also necessitate updating our understanding of neural signatures of temporal prediction. While these signals may originate in extracerebellar circuits, the ability to fine-tune the timing of these neural dynamics is dependent on cerebellar representations in the absence of a periodic context. When coupled with previous work on temporal judgment, reproduction, and sensorimotor learning [63,65,75], we begin to have a coherent picture of the unique role of the cerebellum in temporal attention.

## Methods

### Participants

Eighteen patients with cerebellar degeneration (CD) and 16 neurotypical control individuals were recruited for the study. The data from two individuals from each group were discarded due to excessive noise in the EEG recordings or an inability to perform the task, leading to a final sample size of 16 CD and 14 controls. The study was approved by the Institutional Review Board at the University of California, Berkeley and all of the participants provided informed consent. They were financially compensated for their participation.

Participants in the CD group (10 females, 15 right-handed, mean age=56.7 years, sd=11.4) had been diagnosed with spinocerebellar ataxia, a slowly progressive adult-onset degenerative disorder in which the primary pathology involves atrophy of cells within the cerebellum. We did not test patients who presented symptoms of multisystem atrophy. 11 individuals in the CD group had a specific genetic subtype (SCA1=1, SCA3=5, SCA5=1, SCA6=2, SCA8=1, SCA10=1) and the other five individuals had CD of unknown/idiopathic etiology. All of the CD participants completed a medical history interview to verify the absence of other neurological conditions, and were evaluated at the time of testing with the Scale for the Assessment and Rating of Ataxia (SARA) [76]. The mean SARA score was 11.8 (range: 3.5 – 25.5, sd=6.6). Control participants (8 females, 13 right-handed, mean age=60.4, sd=9.2) were recruited from the same age range as the CD group, and, based on self-reports, did not have a history of neurological or psychiatric disorders. The CD and control groups did not differ significantly in age (*p* = 0.34).

All participants were prescreened for normal or corrected-to-normal vision, intact color vision, and no professional musical training or recent amateur participation in musical activities (e.g., playing a musical instrument or singing in a choir). All of the participants completed the Montreal Cognitive Assessment (MoCA) as a simple assessment of overall cognitive competence. Although we did not select participants to provide a match on this measure, there was no significant group difference (CD: mean = 27.6, Control: mean = 28.3, *p* = 0.18).

### Procedure

Upon arrival, all participants provided consent, demographic information, and completed the MoCA. The CD participants also provided their clinical history (if not on file from a visit within the past year) and were evaluated with the SARA.

The experiment was conducted in a quiet, dimly lit room, with a laptop computer placed on a table in front of the participant. The stimuli (colored squares, 5 x 5 cm, 5.5°) were presented at the center of a 15-in laptop monitor on gray background (viewing distance ≈ 50 cm). Stimulus presentation and response acquisition were handled using Psychophysics toolbox [77] for MATLAB (Mathworks).

Each trial began with the temporal cueing phase, consisting of the serial presentation of two or three red squares (stimulus duration = 100 ms). There were two types of cues, tested in separate blocks. In the Interval task, the cue consisted of two red squares, each presented for 100 ms, with a stimulus onset asynchrony (SOA) of either 700 ms (short cue) or 1200 ms (long cue). In the Rhythm task, the cue consisted of three red squares, presented periodically with an SOA of 700 ms (short cue, equivalent to 1.43 Hz) or 1200 ms (long cue, 0.83 Hz). The last red square was followed by the presentation of a white square, the warning signal (WS), indicating to the participant that the subsequent stimulus would be the target. For the Rhythm task, the interval between the last red square and WS was set to the same duration as the cue SOA for that trial. Thus, the WS fell on the “beat” established by the temporal cues. In contrast, for the Interval task, the interval between the last red square and WS was randomly set on each trial to be either 1.5 or 2.5 times the duration of the SOA on that trial (short cue SOA: 1050 / 1750 ms; long cue SOA: 1800 / 3000 ms). This strongly reduced any periodicity in the stimulus train, and thus eliminated the use of a rhythmic strategy since the WS occurred at 180° phase relative to a “beat” that, in theory, could have been created by the two red squares (see [6]). We favored this type of cuing over purely symbolic cuing (e.g. [8,9]) to minimize the need to learn the target intervals across trials.

Starting with the onset of the WS, the trial events were identical for the two tasks. After a short delay, the target, a green square was presented, and the participant was instructed to make a speeded button press using their right index finger upon detection of the target. The interval between the WS and target was either the same SOA as defined by the temporal cue (valid trial) or the non-cued SOA (invalid trials). The cue was valid on 56.25% of the trials and invalid on 18.75% of the trials; this 3:1 ratio was selected to incentivize the participant to attend to the temporal cues to facilitate performance. On the remaining 25% of the trials, no target was presented. These catch trials were included to discourage participants from making anticipatory responses [10]. Participants received feedback (error message on the monitor) if the responded prematurely, responded on a catch trial, or if they did not respond within 3 s of target onset.

Participants preformed 4 blocks of each task, each consisting of 32 trials, in alternating order (8 blocks total, first block counter-balanced across participants). Within each block, the duration of the temporal cue was randomly determined with the constraint that each cue occurred on 50% of the trials for all conditions (valid, invalid, catch). Short breaks were provided between each block. Prior to the first block for each task, the experimenter demonstrated the trial sequence and then conducted short blocks of practice trials (n=8 trials), repeating the practice block until the participant could describe how the cues were predictive of the onset time of the target. For subsequent blocks, the participant first completed two practice trials as a reminder of the format for the temporal cues in the forthcoming block.

### EEG recording and preprocessing

EEG was recorded continuously from 64 preamplified Ag/AgCl electrodes, using an Active 2 system (BioSemi, The Netherlands). The electrodes were mounted on an elastic cap according to the extended 10–20 system. Additional electrodes were placed on the outer canthi of the right and left eyes, and above and below the center of the right eye to track electro-ocular activity, and on the left and right mastoids and near the tip of the nose to be used as reference electrodes. The EEG signal was sampled at a rate of 1024 Hz (24 bits/channel), with an online anti-aliasing 204 Hz low-pass filter.

EEG preprocessing was conducted in MATLAB using the FieldTrip toolbox and custom written scripts. We employed the following analysis pipeline: 1) Referencing to average of right and left mastoid electrodes. 2) High-pass filtering using a zero-shift Butterworth filter with a cutoff of 0.1 Hz (24 dB/octave). 3) Correction of ocular artifacts using independent component analysis (ICA, [78]) based on typical scalp topography and time course. 4) Elimination of epochs which contained artifacts that were non-cognitive in origin (defined as absolute activity larger than 100 μV, or a change of more than 100 μV in a 200-ms interval).

### Behavioral data analysis

Trials were discarded if a response was detected before target onset, or if the RT was shorter than 100 ms or longer than 3000 ms (2.2% of trials, no difference between groups or tasks). From the remaining trials, trials were discarded if the RT was more than three standard deviations above or below the mean RT, calculated separately for each of the conditions. Only 0.7% of all trials were rejected on this criterion, with no difference between groups or tasks. The RT data of each task were then subjected to a mixed ANOVA with factors Group (CD / Control), Target SOA (early / late), and Cue Validity (valid / invalid). To assess the validity effect within each group we conducted planned contrasts on the cue validity factor, subjecting the data of each group to a repeated-measures ANOVA with factors Cue Validity (valid / invalid) and Target SOA (short / long). To assess context-specificity within the CD group, we performed a secondary analysis with the orthogonal contrast, using a repeated-measures ANOVA with factors Task (Interval/Rhythm), Target SOA, and Cue Validity. Here and in all subsequent analyses, effect sizes were estimated using Cohen’s d and partial eta-squared 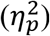.

### ERP analysis

All EEG analyses were conducted in MATLAB using custom written scripts and the CircStat toolbox [79]. Analysis of anticipatory ERP components focused on the CNV, a negative polarity potential arising in the interval between the WS and an expected target. This response typically peaks just prior to the anticipated event [6–8]. For the CNV analysis, continuous EEG data was segmented into epochs extending from 200 ms before to 700 ms after WS stimuli, and these were averaged separately for each participant, task and expected SOA. The 100 ms epoch just before the WS was used as the baseline. Given that the early target would occur 700 ms after the WS, we predicted that the CNV would have a larger amplitude (i.e., be more negative) following the short temporal cue, relative to the long temporal cue.

Analysis of a target-evoked ERP focused on the P3 response, whose amplitude is typically enhanced following valid compared to invalid temporal cues [8,45]. Note that while temporal prediction were also found to impact early sensory EEG responses, these effects are only observed in demanding perceptual tasks [49]; they are typically absent in supra-threshold detection tasks such as that employed here. For the P3 analysis, segments extending from 200 ms before to 700 ms after target onset were averaged separately for each participant, task, and cue validity, with a period of 100 ms before the target used as baseline. Note that we only analyzed the P3 to short SOA targets. This avoids baseline artifacts that might occur for long SOA targets on invalid trials due to changes in the CNV following the absence of the expected target at the short SOA [6,7].

The CNV and P3 were analyzed in pre-defined fronto-central (Fz, FC1, FCz, FC2, Cz) and parietal (P1, Pz, P2) electrode clusters, respectively [6–8]. Electrode selections were validated by inspecting the data across groups and conditions (Figure 2A, 5A). To compare the CNV and P3 amplitude between conditions, the amplitude was averaged for each participant across a predefined time window (CNV: 500-700 ms after the W S, just prior to the short SOA target; P3: 275-325 ms after the target, around the expected peak latency based on the literature and an informal assessment of our data set across conditions). For each task, these values were compared within each group using a paired t-test, and the modulation of each ERP by the cue duration was compared between groups using a mixed ANOVA. As with the RT data, we assessed context-specificity in the CD group using a repeated-measures ANOVA with factors Task (Interval/Rhythm) and Cue SOA.

We also used a using a cluster-based permutation test to evaluate the temporal extent of the difference between conditions without restriction to a pre-defined window [40] (10000 iterations, shuffling conditions within participants, individual time point threshold: *p* < 0.05).

Randomization test p-value can vary slightly with repeated application of the analysis due to the arbitrary shuffling. As such, here and in subsequent permutation analyses p-values are reported as lower than a significance threshold.

To quantify how the CNV was affected by cerebellar degeneration in each task, we fit the CNV waveform observed between the WS and the time of the short SOA target (700 ms) with a two-parameter model of ramping activity (Figure 2D). The model approximates a climbing neuronal activity process [80,81], assumed to have linear ramping (slope, *S*) with a variable onset time (*T*) as follows:

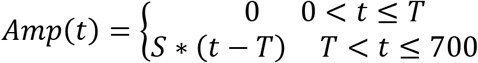

This model was fit to the data of each task and cue duration (averaged across participants) using the “fminsearch” function in MATLAB (Nelder–Mead simplex optimization) with several initial values to minimize effects of local minima. To compare the model parameters, we used a randomization approach in which we first calculated the difference between parameters in the original data (e.g., slope difference between short and long cues). We compared these values to a null distribution of the difference between parameters, obtained by fitting the model to randomized datasets (formed by shuffling condition labels within participants, 5000 iterations). For each parameter, the difference between conditions (group or cue duration) was considered significant only if it was larger than the difference obtained in 95% of the randomized datasets.

### Time Frequency Analysis

For the time frequency analysis, we focused on modulations in the delta and beta bands. In the delta band, temporal prediction is associated with increased inter-trial phase concentration (ITPC, [42]) in anticipation of a target [3,4,6], in both periodic and aperiodic streams [6,35,36]. For the ITPC analysis, we band-pass filtered the data in the delta-frequency range (0.6-2 Hz Butterworth filter, 24 dB/octave), extracted the instantaneous phase using the Hilbert transform, and calculated the ITPC (Figure S2). We chose to use the 0.6 to 2 Hz frequency range given that it is symmetric (logarithmically) around the frequencies that correspond to the short and long target intervals (700 ms = 1.43 Hz and 1200 = 0.83 Hz; ~0.5 octave margins from each side). We used a causal filter instead of a non-causal zero-lag filter to avoid contamination of the anticipatory period by target-evoked activity.

To examine the effect of cerebellar degeneration on ITPC modulation by temporal prediction, we applied this analysis pipeline to the EEG data from the same fronto-central ROI as used in the CNV analysis (Fz, FC1, FCz, FC2, Cz). The data were segmented within an epoch extending from 200 ms prior to the WS until 700 ms after the WS, with ITPC calculated at each time point, separately for each group, task and cue duration. Given that we did not expect to observe a difference in this measure between the two temporal cue conditions, we averaged the results across this factor. We focused on ITPC levels just prior to the early target by averaging across a pre-defined window of 600-700 ms after the WS. These values were baseline corrected, and then compared between groups. For the baseline, ITPC values were averaged across the 100 ms epoch just prior to the onset of the WS. The pre-WS period from the Interval task was used as baseline for both tasks, given that, in the Rhythm task, the WS is also temporally predictable. Since ITPC values are not normally distributed, the groups were compared using a non-parametric permutation test. We compared the observed ITPC difference to a null distribution of randomized ITPC differences, created by shuffling group labels (5000 iterations). Context-specificity in the CD group was tested with a non-parametric permutation test in which the null distribution was created by shuffling task labels.

To examine whether ITPC, when calculated in this manner, can reflect temporal consistency of non-oscillatory ramping activity, we applied our pipeline to simulated data. We simulated a set of 100 trials (matching the approximate number of trials per condition in the actual data) using the 2-parameter ramping activity model described above. To avoid edge artifacts, we added a stage in which the activity gradually returns to baseline after target onset [6,7]. For each set of simulated trials, ramping onset latency and slope for each trial were chosen randomly from a Gaussian distribution with a specific standard deviation (Onset latency: mean = 300 ms, standard deviation= 0, 50, 100, 150 or 200 ms; Slope: mean = −12 μV/sec, standard deviation = 0, 5, 10, 15, 20 μV/sec). 1/f random noise was added to the simulated time series given its presence in EEG recordings. We then applied the ITPC pipeline used for the EEG data and extracted the ITPC just prior to target time. To ensure stability, the results for each of the 25 combinations of distribution standard deviations were averaged across 100 iterations and three amplitude levels of the 1/f noise. To examine whether ITPC depended on the magnitude of inter-trial variability in ramping onset latency or slope, we conducted a multiple regression analysis to predict the resulting ITPC values from these two predictors (see also Figure S2).

The amplitude of activity in the beta band decreases prior to the time of an expected event that requires a motor response [7,15,17,82]. Thus, we expected lower beta amplitude just prior to the time of the early target following a short cue compared to following a long cue. To test this prediction, segments extending from 1200 ms before to 2200 ms after the WS were subjected to a time-frequency decomposition using a complex Morlet wavelet transform (1-40 Hz, 1 Hz steps, ratio between the central frequency and the standard deviation of the Gaussian-shaped wavelet in the frequency domain = 8). We discarded the first and last 1,000 ms from the analyses to exclude edge artifacts. Instantaneous amplitudes were averaged separately across trials for each participant, task and temporal cue for each electrode. For the baseline measure of spectral amplitude, we used the 100 ms window after the WS (see [6]). In measuring the change in spectral amplitude, we only included trials in which target did not appear at the early interval (invalid and catch for short cue, valid and catch for long cue) to avoid contamination from target-evoked activity.

Beta-band suppression is observed in central-parietal sites, but there is no consensus on the appropriate frequency range for beta. To select the range for our analyses in an unbiased manner, we inspected the time-frequency representations, averaged across groups, tasks and electrodes (Figure S3). From this grand average function, we identified suppression in the frequency range of 14-26 Hz. Examining the scalp distribution of activity in these frequencies confirmed a central-parietal focus. Therefore, we analyzed beta activity in a central-parietal cluster (C3, C1, Cz, C2, C4, CP3, CP1, CPz, CP2, CP4).

To compare the beta amplitude between conditions, the amplitude was averaged for each participant across a pre-defined time window (500-700 ms after the WS, just prior to the early target time). For each task, a within group comparison of the short and long cue conditions (restricted to the data from trials in which the target appeared at the long SOA) was conducted using a paired t-test. For the between group analyses, we used a mixed ANOVA (Group x Cue SOA). We also evaluated the spectro-temporal extent of the difference between conditions without commitment to a pre-defined time or frequency window using a two-dimensional cluster-based permutation test [40] (10000 iterations, shuffling conditions within participants, individual time point threshold: *p* < 0.05). Context-specificity in the CD group was evaluated with a repeated-measures ANOVA with factors Task (Interval/Rhythm) and Cue SOA.

## Acknowledgements

We are grateful to Katherine Duberg, Claudia Tischler, and Katie Andrade for their assistance in collecting the data. This work was supported by grants from the National Institute of Health (NS092079, NS105839).

## Author Contributions

Conceptualization, A.B. and R.B.I.; Methodology, A.B. and R.B.I.; Software, A.B.; Validation, A.B. and R.B.I.; Formal Analysis, A.B.; Investigation, A.B.; Data Curation, A.B. and R.B.I.; Writing – Original Draft, A.B.; Writing – Review & Editing, A.B. and R.B.I.; Visualization, A.B. and R.B.I.; Supervision, R.B.I; Funding Acquisition, R.B.I.

## Supplementary figures

**Figure S1.**
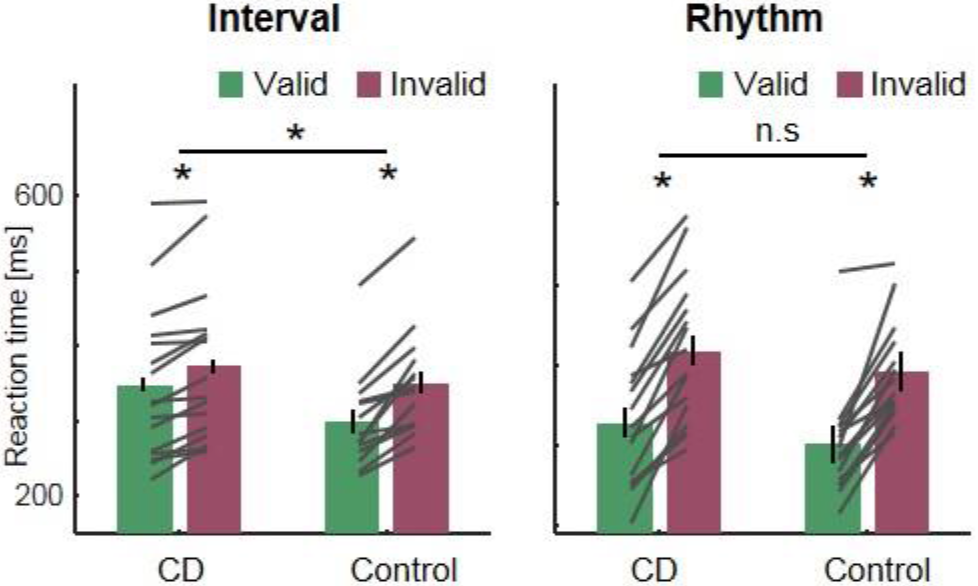
Behavioral results with single-subject data. Same conventions as in Figure 1C, D.

**Figure S2.**
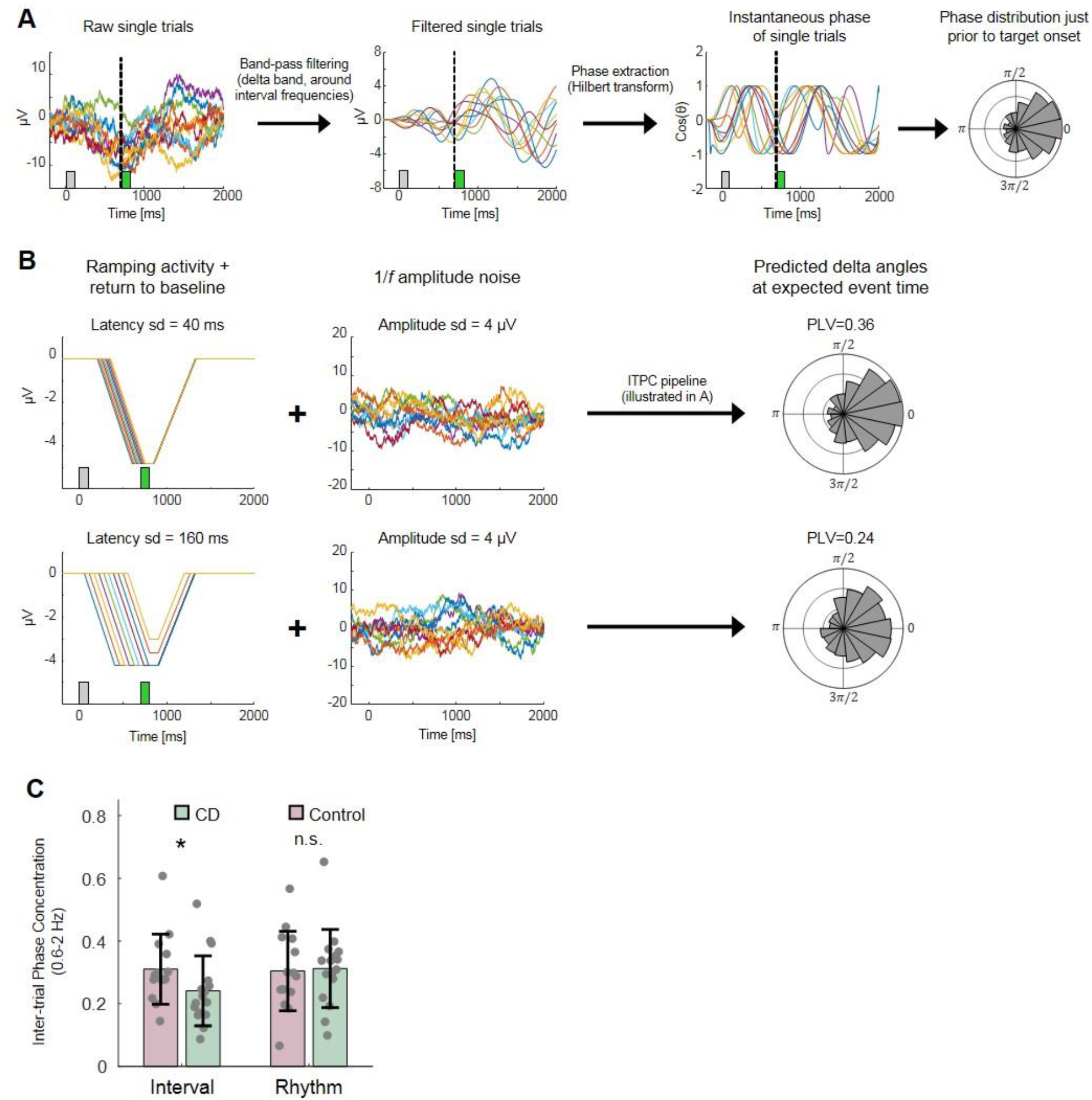
Delta-band analysis. A. Analysis pipeline. The data were filtered using a 0.6-2 Hz band-pass causal filter, and the Hilbert transform was applied to obtain instantaneous phase. The intertrial phase concentration (ITPC) was calculated across trials for each time point (the depicted phase distribution is from the last time point prior to target onset, marked with dashed line). B. Simulated trials were created using the ramping model (see ERP analysis), and then adding random-walk 1/f noise. Phase distributions are demonstrated for the 700 ms time point, corresponding to when the early target would appear (green square). C. ITPC values measured in the EEG data from the CD and control groups for the interval and rhythm tasks, with individual subject data. Error bars represent one standard error of the mean (SEM) difference between tasks. * p < 0.05.

**Figure S3.**
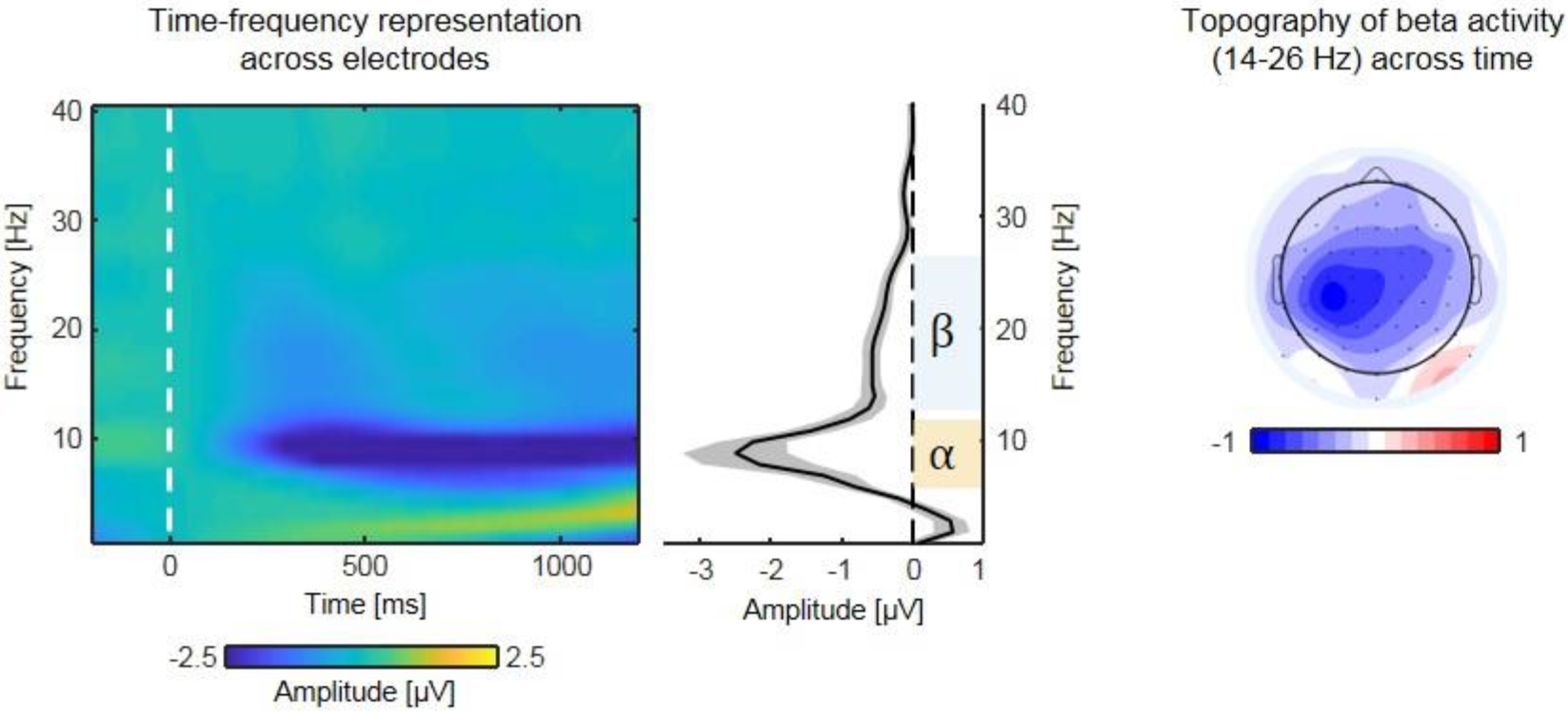
Defining the beta-band frequency range of interest. Left: baseline corrected time-frequency representation across electrodes, groups, tasks and expected SOAs. The data are from trials in which the target appeared at the long SOA or from catch trials. Oscillatory suppression is observed in the alpha and beta bands. Middle: spectral density across time, showing beta-band suppression (significantly lower than 0) in 14-26 Hz. Error margins represent one standard error of the mean (SEM) difference from baseline (uncorrected for multiple comparisons). Right: Scalp distribution of activity averaged across the identified beta range, across groups and tasks, shows a typical central-parietal distribution.

